# The novel roles of choline transporter-like 1 and 2 in ethanolamine transport

**DOI:** 10.1101/2020.08.27.270223

**Authors:** Adrian Taylor, Sophie Grapentine, Jasmine Ichhpuniani, Marica Bakovic

## Abstract

We examined a novel function of mammalian Choline-Transporter-Like proteins CTL1/SLC44A1 and CTL2/SLC44A2 in ethanolamine transport. We established two distinct ethanolamine transport systems of a high affinity (*K*_*1*_ = 55.6 - 66.5 μM), mediated by CTL1, and of a low affinity (*K*_*2*_ = 275 - 299 μM), mediated by CTL2. Both types of transport are Na^+^-independent and mediated in a pH dependent manner, as expected for ethanolamine/H^+^ antiporters. Primary human fibroblasts with separate frameshift mutations (M1= *SLC44A1* ^ΔAsp517^ and M2= *SLC44A1* ^ΔSer126^) are devoid of CTL1 ethanolamine transport but maintain unaffected CTL2 transport. The lack of CTL1 or CTL2 reduced the ethanolamine transport, the flux by the CDP-ethanolamine Kennedy pathway and PE synthesis. Overexpression of CTL1 in *SLC44A1* ^ΔSer126^ (M2) cells improved the ethanolamine transport and PE synthesis. The *SLC44A1* ^ΔSer126^ cells are reliant on CTL2 function and CTL2 siRNA almost completely abolished ethanolamine transport in the whole cells and mitochondria. Overexpression of CTL1 and CTL2 cDNAs increased ethanolamine transport in control and *SLC44A1*^ΔSer126^ cells. CTL1 and CTL2 facilitated mitochondrial ethanolamine uptake, but the transport mediated by CTL1 is predominant in the whole cells and mitochondria. These data firmly established that CTL1 and CTL2 are the first identified ethanolamine transporters in the whole cells and mitochondria, with intrinsic roles in *de novo* PE synthesis by the CDP-Etn Kennedy pathway and compartmentation of intracellular ethanolamine.

**Significance:** The lack of Choline Transporter Like 1 (*SLC44A1*/CTL1) is the primary cause of a new neurodegenerative disorder with elements of childhood-onset parkinsonism and mitochondrial dysfunction. *SLC44A2*/CTL2 encodes the human neutrophil antigen 3, causes autoimmune hearing loss and Meniere’s disease, and has been recently identified as the main risk factor for thrombosis-the major cause of death in Covid-19 patients. Our investigation provides insights into the novel functions of CTL1 and CTL2 as intrinsic ethanolamine transporters. CTL1 and CTL2 are high and low affinity transporters, with direct roles in the membrane phospholipid synthesis. The work contributes to new knowledge for CTL1 and CTL2 independent transport functions and the optimization of prevention and treatment strategies in those various diseases.

## Introduction

Phosphatidylcholine (PC) and phosphatidylethanolamine (PE) are major components of cellular membranes where they are involved with essential cellular processes (1, 2). PC and PE are synthesized *de novo* by CDP-Cho and CDP-Etn branches of the Kennedy pathway in which the extracellular substrates choline (Cho) and ethanolamine (Etn) are actively transported into the cell, phosphorylated and coupled with diacylglycerols (DAG) to form the final phospholipid product. While multiple transport systems have been established for Cho, Etn transport is poorly characterized and there is no single gene/protein assigned a transport function for mammalian Etn. Cho transport for membrane phospholipid synthesis is mediated by Cho transporter like protein CTL1/SLC44A1 (3). CTL1 is the only well-characterized member of a broader family (CTL1-5/SLC44A1-5) (4, 5). CTL1/SLC44A1 is a Cho/H^+^ antiporter at the plasma membrane and mitochondria (4, 5). The role of plasma membrane CTL1 is assigned to Cho transport for PC synthesis, but the exact function of the mitochondrial CTL1 is still not clear. In the liver and kidney, mitochondrial CTL1 transports Cho for oxidation to betaine, the major methyl donor in the one-carbon cycle (6). In other tissues however, the mitochondrial CTL1 probably maintains the intracellular pools of Cho and as a H^+^-antiporter and modulates the electrochemical/proton gradient in the mitochondria (7, 8). CTL2/SLC44A2 is only indirectly implicated in PC synthesis and its exact function is not firmly established in neither whole cells nor mitochondria (4).

PE is the major inner membrane phospholipid with specific roles in mitochondrial fusion, autophagy and apoptosis (9 - 11). PE is also a valuable source of other phospholipids. PC is produced by methylation of PE while phosphatidylserine (PS) is produced by an exchange mechanism whereby the Etn moiety of PE is replaced with serine and free Etn is released. PC could also produce PS by a similar exchange mechanism, with free Cho being released. The metabolically released Cho and Etn need to be transported in and out of the cytosol and mitochondria or reincorporated into the Kennedy pathway (3 - 6). That mammalian Etn and Cho transport may occur through a similar transport system was implicated from early kinetic studies in bovine endothelial cells, human retinoblastoma cells and glial cells (12 - 14). Here, we demonstrate that CTL1/SLC44A1 and CTL2/SLC44A2 are authentic Etn transporters at the cell surface and mitochondria. We examine the kinetics of Etn transport in CTL1 and CTL2 depleted conditions and overexpressing cells. We characterize Etn transport in human skin fibroblasts that maintain CTL2 but lack CTL1 function due to inherited *CTL1/SLC44A1* frameshift mutations (M1= *SLC44A1* ^ΔAsp517^ and M2= *SLC44A1* ^ΔSer126^) (15). We employ pharmacological and antibody induced inhibition to separate the contributions of the CTL1 and CTL2 to Etn transport and PE synthesis. This study is the first to demonstrate that the CTL1 and CTL2 are high and low to medium affinity cellular and mitochondrial Etn transporters. To our knowledge, this is the first study to demonstrate that as intrinsic Etn transporters, CTL1 and CTL2 regulate the supply of extracellular Etn for the CDP-Etn pathway, redistribute intracellular Etn and balance CDP-Cho and CDP-Etn arms of the Kennedy pathway.

## Results

### CTL1 and CTL2 inhibition reduces Etn and Cho transport

To assess the magnitude by which CTL1/2 inhibition affects PE and PC levels, two types of cells were characterized for phospholipid metabolism (MCF-7 and MCF-10) (16, 17). The cells were treated for 24h with [^3^H]-glycerol, to label the entire glycerolipid pools (steady-state levels) in the presence and absence of CTL1 transport inhibitor hemicholinium-3 (HC-3) or CTL1 specific antibody (Fig. 1A and B). Surprisingly, HC-3 reduced the steady-state levels not only of PC (25-50%) but also of PE (50%) in both cell types. CTL1 antibody similarly reduced PC and PE levels (40-50%), further indicating that CTL1 could be involved in the transport of Etn, in addition to its well-characterized function in Cho transport (15, 18). [^3^H]-Cho and [^14^C]-Etn transport were similarly inhibited with the CTL1 inhibitor HC-3 (Fig. 1C) and they compete for the same transport system (Fig. 1D). [^14^C]-Etn and [^3^H]-Cho transport were similarly diminished when equal, 200 μM, Etn, Cho and Etn + Cho were applied respectively (Fig. 1D). Furthermore, CTL1 antibody inhibited ^14^C-Etn transport in a concentration-dependent manner with an IC_50_ of 50 ng (Fig. 1E). CTL2 antibody also inhibited both, ^14^C-Etn and ^3^H-Cho transports in a concentration dependent manner with LC_50_ 50 ng (Fig. 1F, G). Together, the data showed that Etn is a 3ubstrate for CTL1 and CTL2-mediated transports, in addition to already establish functions in Cho transport.

**Figure 1:**
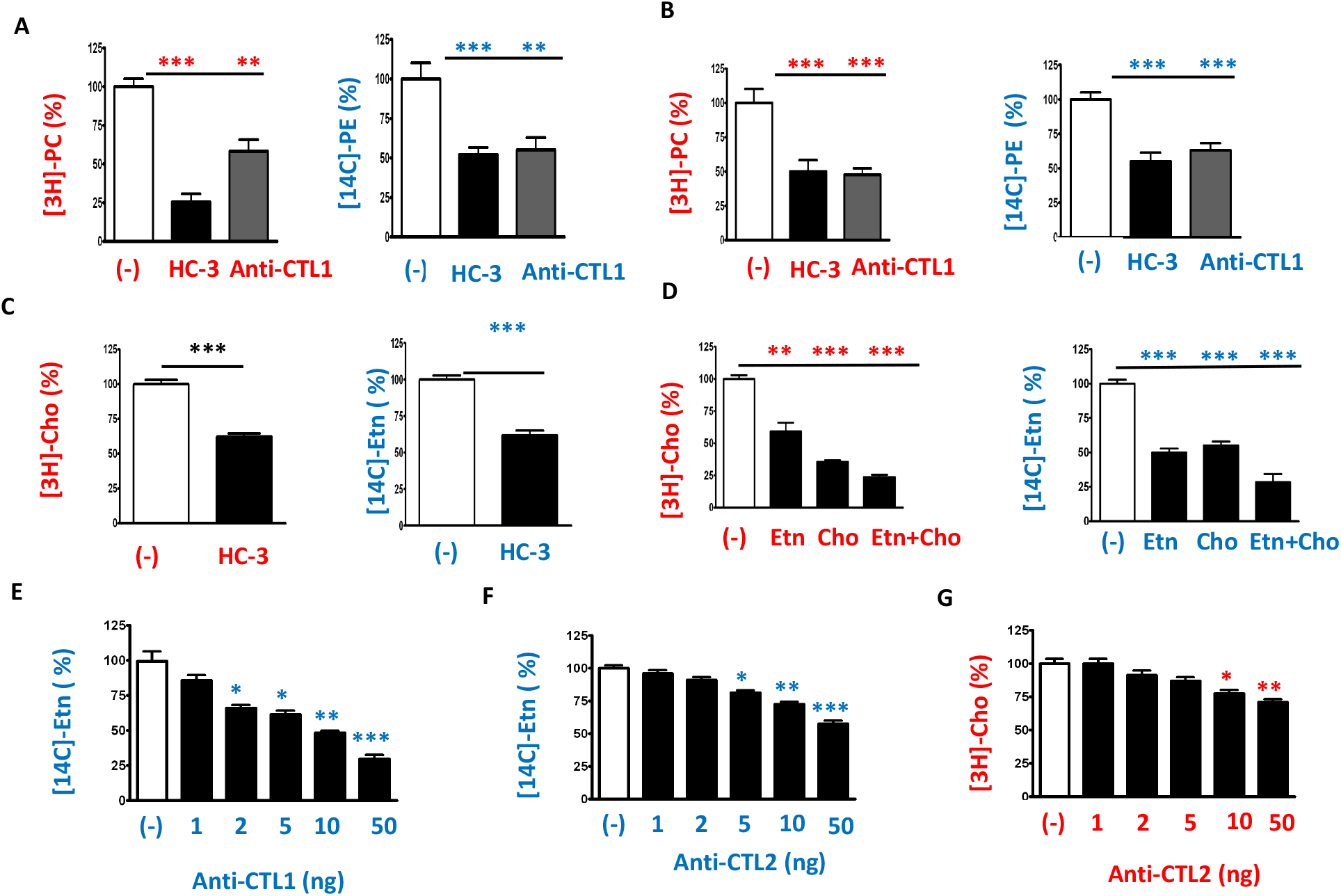
Ethanolamine transport resembles CTL1- and CTL2-mediated choline transport. **(A, B)** MCF-7 (A) and MCF-10 (B) cells were pretreated with CTL1/2 inhibitor hemicholinium (HC-3, 200μM) or anti-CTL1 antibody (1:500) and then radiolabeled with 0.2 μCi [^14^C]-Etn or [^3^H]-Cho for 24h. Both [^3^H]PC and [^14^C]PE were significantly reduced in HC-3 and anti-CTL1 treated cells relative to untreated (-) cells. **(C)** 0.2 μCi [^14^C]-Etn or [^3^H]-Cho uptake measured after 20 min in MCF7 cells were significantly reduced with HC-3 (200 μM) added 20 min prior to the labeling). **(D)** Separately or together, excess of ‘cold’ Cho and Etn (200 μM each, 20 min) inhibited the transport of both radiolabeled substrates. **(E-G)** Anti-CTL1 and anti-CTL2 antibodies inhibited [^14^C]-Etn (E, F) and [^3^H]-Cho (G) transport in primary human fibroblasts in a dose-response manner. Each bar represents the mean ± SD (n = 4); * p < 0.05, ** p < 0.01, *** p < 0.001.

### CTL1 is a high-affinity and CTL2 is a low-affinity Etn transporter

We studied the kinetics of Etn transport in monkey COS-7 cells and control (Ctrl) and CTL1 deficient (M1=*SLC44A1*^ΔAsp517^ and M2=*SLC44A1*^Ser126^) primary human fibroblasts. As expected, COS-7 cells and Ctrl fibroblasts expressed CTL1 and CTL2 proteins while CTL1 mutant fibroblasts M1 and M2 only expressed CTL2 protein (Fig. 2A). ^14^C-Etn transport rates (V) plotted against [Etn] produced a series of saturation curves, as expected for protein mediated transports (Fig. 2B). *V*_*max*_ values were nearly identical in Ctrl and COS-7 cells (*V*_*max*_ *=* 26.9 and 26.3 nmol/mg protein/min) and M1 and M2 cells had reduced but similar *V*_*max*_ = 20.6 - 21.2 nmol/mg protein/min (Fig. 2B), apparently caused by the absence of the CTL1 transport component. Indeed, the Eadie-Hofstee plots derived from the saturation curves were biphasic in Ctrl fibroblasts and COS-7 cells and linear for M1 and M2 cells (Fig. 2C). This type of behavior indicated the presence of two distinct transport systems in Ctrl and COS-7 cells with two binding constants, of high and low affinity for Etn, and one transport system of a lower affinity in M1 and M2 cells. As further shown in Fig. 2C, Ctrl fibroblasts, high affinity (K_1 =_ 66.5 ± 8.5 μM) and low (K_2_ = 299.0 ± 13.1 μM) affinity Etn bindings were similar to COS-7 cells bindings (K_1_ = 55.6 ± 14.8 μM and K_2_ = 277.3 ± 7.9 μM). On the other hand, M1 and M2 cells are characterized by a single transport with a binding constant for Etn of 275.4 – 279.6 μM which is the second (K_2_), low affinity, binding constant as determined in Ctrl and COS-7 cells (Fig. 2C). M1 and M2 cells only express CTL2 and at levels similar to Ctrl and Cos 7 cells, and do not have a functional CTL1 protein (Fig. 2A,D), strongly implicating CTL2 as responsible for the low affinity Etn transport. Indeed, CTL2 depletion by siRNA knockdown in Ctrl cells completely abolished the low affinity transport component while the high affinity component remained intact (Fig. 2E). This analysis also confirmed that the high affinity transport (K_1_) which is absent in M1 and M2 cells and remained intact in CTL2 siRNA treated Ctrl is CTL1-mediated Etn transport.

**Figure 2:**
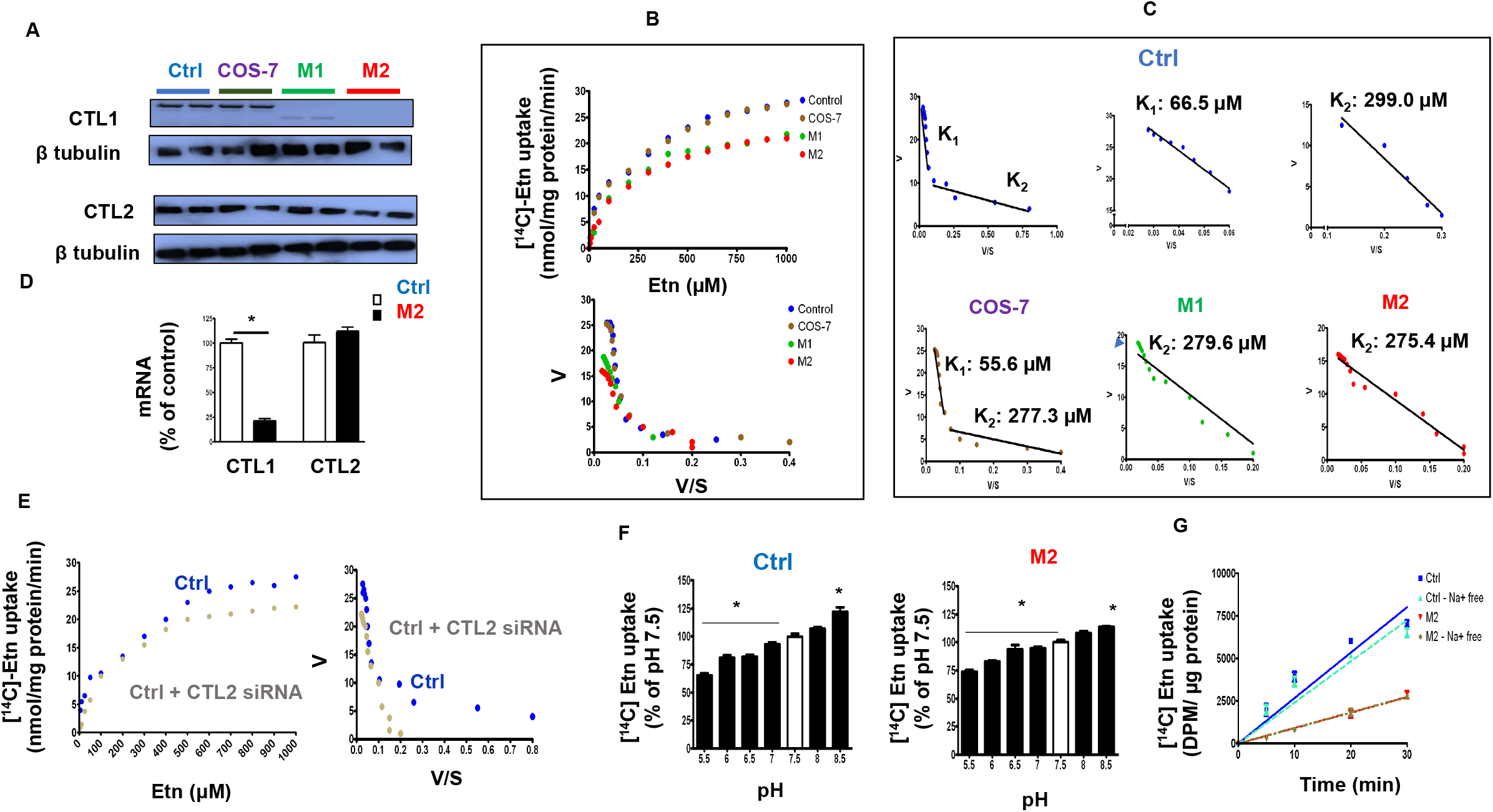
Characteristics of low and high affinity Etn transports. **(A**) Saturation curves of Etn transport (V (velocity) vs. S (substrate conc), were produced by measuring the uptake of [^14^C]-Etn (0-1000 μM, 20 min) in control human fibroblasts (Ctrl), CTL1 deficient human fibroblasts (M1 and M2) and monkey COS-7 cell line **(B)** The Eadie-Hofstee plots derived from those curves demonstrated the presence of two Etn transport systems, with binding constants K1 (higher affinity) and K2 (lower affinity), in Ctrl and COS-7 cells. Only one transport system with the lower affinity was present in CTL1 deficient M1 (279.6 ± 17.1 μM (K_2_)) and M2 (275.4 ± 13.9 μM (K_2_)) cells. **(C)** CTL1 protein (72 kDa) was detected in Ctrl and COS-7 cells while a truncated, low abundant 58 kDa protein was detected in M1 cells. No CTL1 protein was detected in M2 cells showing that the Eadie-Hofstee plots K2 Etn transport system in those cells is CTL2 related. An intact 72 kDa CTL2 protein was detected in M1 and M2 cells. **(D)** CTL2 was not affected while CTL1 mRNA was almost diminished in M2 cells. **(E)** The [^14^C]-Etn uptake is down-regulated with low extracellular pH (high [H^+^]) and upregulated at high pH (low [H^+^]) in both Ctrl (CTL1+CTL2) and M2 (CTL2 only) cells. **(F)** Saturation curves and Eadie-Hofstee plots of siRNA-CTL2 treated Ctrl cells showed the presence of a single Etn transport of a higher affinity (CTL1-mediated, K1, transport); the low affinity, CTL2-mediated, K2 transport was specifically depleted with the siRNA-CTL2 treatment. **(G)** Time course (0-30 min) of 10 μM [^14^C]-Etn uptake in the presence and absence of Na^+^ ions in Ctrl and M2 cells. Each bar or point represents the mean ± SD (n = 4), * p < 0.05, ** p < 0.01.

Since the effects of pH and [Na^+^] ions on choline transport is well established (19), their effects on Etn transport were also investigated (Fig. 2F and G). Etn transport in Ctrl (CTL1 + CTL2 transport) and M2 fibroblasts (CTL2 transport) (Fig. 2F) was reduced when extracellular pH was lowered from 7 to 5.5 and stimulated when pH was increased to 8.5. Additionally (Fig. 2G), as expected, the rate of Etn transport in Ctrl cells was higher than in M2 cells but the rates were not modified when Na^+^ ions were replaced by Li^+^ ions in the uptake buffer. Altogether, the data established that CTL1 and CTL2 acts as Etn/H^+^ antiporters, driven by a proton gradient and they are both independent of Na^+^, as in case of Cho transport (19).

### CDP-Etn Kennedy pathway is down regulated by CTL1 deficiency

Since this is the first time that Etn transporters are identified, it is important to establish if they are functionally linked to the CDP-Etn Kennedy pathway for PE synthesis. We performed pulse (synthesis) and pulse-chase (degradation) radiolabeling with [^14^C]Etn in Ctrl and CTL1 deficient M2 cells, to establish the contributions of total and CTL2 transport to the CDP-Etn Kennedy pathway (Fig. 3A,B). As shown in (Fig. 3A), the total incorporation of [^14^C]Etn and the rates of synthesis (slopes) of the pathway intermediates phospho-Etn (P-Etn), CDP-Etn and the final product PE were lower in M2 cells than in Ctrl cells. Similarly reduced was the [^14^C]Etn incorporation and P-Etn and CDP-Etn disappearance in M2 cells in the pulse-chase experiments (Fig. 3B). In addition, the [^3^H]glycerol radiolabeling of glycerolipid equilibrium pools (the steady-state levels) showed unchanged PC, reduced PE, PS and DAG and increased triglycerides (TAG) in M2 cells (Fig. 3C and D). Therefore, reduced Etn transport, slower P-Etn and CDP-Etn formation, and reduced DAG levels, collectively slowed the CDP-Etn pathway (Fig. 3A,B) and reduced PE levels (Fig. 3C) in M2 cells.

**Figure 3:**
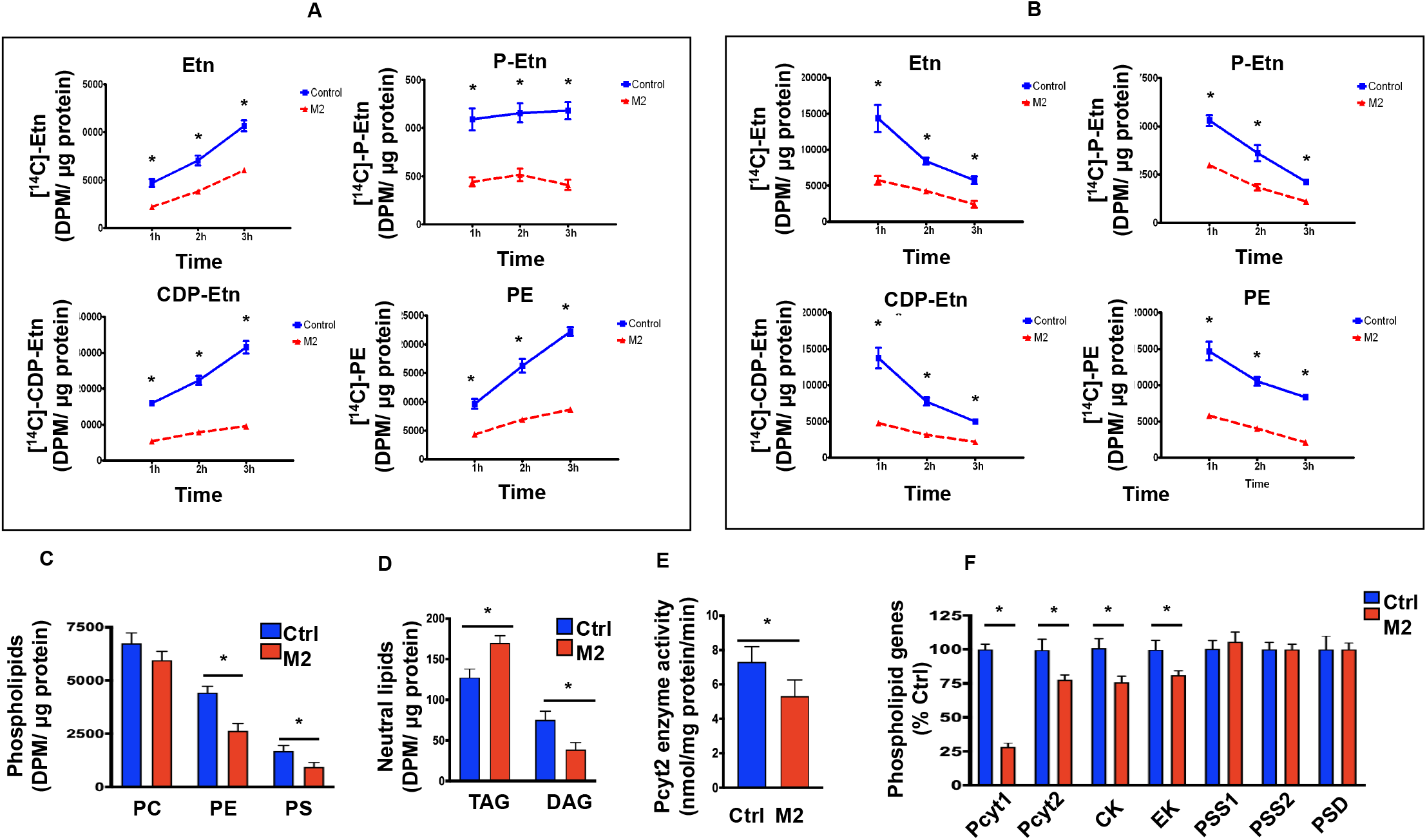
Contributions of CTL2 to PE synthesis by the CDP-Etn Kennedy pathway. **(A**) [^14^C]-Etn 1-3 h radiolabeling of CDP-Etn Kennedy pathway intermediates (Etn, P-Etn and CDP-Etn) and PE synthesis in Ctrl and M2 cells. **(B)** Pulse-chase [^14^C]-Etn labelling of the CDP-Etn pathway intermediates and PE turnover in Ctrl and M2 cells. **(C**,**D)** [3H]Glycerol labelling of glycerolipid pools showing how PC was unchanged, while PE and PS were decreased in M2 cells relative to Ctrl cells (C); and how TAG levels were increased whereas DAG was decreased in M2 cells (D). **(E)** Pcyt2 activity was significantly reduced in M2 cells. **(F)** Pcyt1 mRNA expression was decreased by 65% in M2 cells, while Pcyt2, CK, and EK mRNA expression were decreased by 20% in M2 cells. Each bar or point represents the mean ± SD (n = 4), * p < 0.05.

The CDP-Etn formation from PEtn is usually the rate-regulatory step in the Kennedy pathway and is controlled by Pcyt2 (CTP: phosphoethanolamine cytidylyltransferase) (11). Indeed, in accordance with reduced CDP-Etn formation above, the activity and expression of Pcyt2 were also reduced in M2 cells (Fig. 3 E,F). The expression of Etn kinase (EK) was similarly decreased by 25% (Fig. 3F), explaining why the formation of P-Etn was reduced in M2 cells (Fig. 3A, B). In addition to PE, PS levels were reduced in M2 cells but the expression of PS synthesis (PS syntase1/2-PSS1/2) and PS degradation (PS decarboxylase-PSD) genes (Fig.3F) were unaltered (Fig 3C). The expression of the PC synthesis genes Pcyt1 (CTP: phosphocholine cytidylyltransferase) was decreased by 70% and choline kinase (CK) by 25% in M2 cells (Fig. 3F), yet unexpectedly PC levels were unchanged (Fig. 3C). We previously established (15) that the constant PC levels in CTL1 deficient M1 and M2 cells are maintained by reduced PC turnover and increased formation from other phospholipids (PC is made at the expanse of PE and PS), as the main mechanism to maintain PC as a source of choline in a new neurodegenerative disorder caused by frame-shift mutations in the CTL1 gene M1=*SLC44A1*^ΔAsp517^ and M2=*SLC44A21*^Ser126^ (15). These data collectively provided strong genetic and metabolic evidence that CTL1 and CTL2 independently contributes not only to the CDP-Cho but also to the CDP-Etn Kennedy pathway. CTL2 is not over expressed in deficient M1 and M2 cells (Fig.2A) and as such it cannot compensate for the absence of CTL1 in those cells and affected individuals (15).

### Overexpressed CTL1 and CTL2 participate in choline and ethanolamine transport

To demonstrate that CTL1 and CTL2 are both Etn and Cho transporters, the cells were transiently transfected with CTL1 cDNA or CTL2 cDNA and the protein expression and transport determined after 48h. As shown in Fig. 4A, in M2 cells, that completely lack CTL1 protein, CTL1 cDNA increased CTL1 protein to the levels in Ctrl cells. In Ctrl cells, that have abundant CTL1 and CTL2, transfections only modestly increased the protein levels. On the other hand, when cells were treated with CTL2 siRNA the treatment almost completely abolished CTL2 protein in both cells, which diminished the low affinity Etn transport, described in Fig 2E.

**Figure 4:**
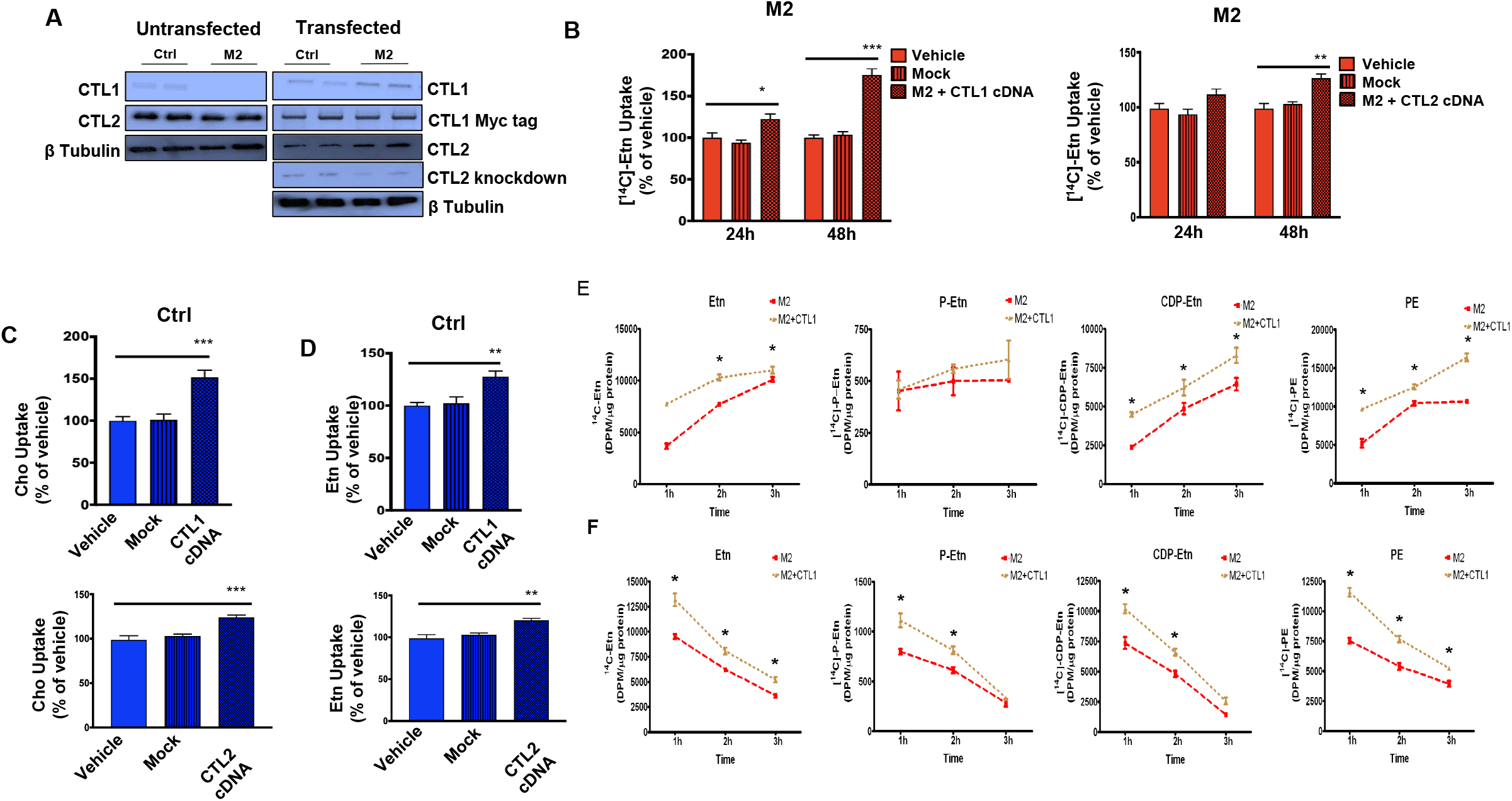
CDP-Etn Kennedy pathway is regulated with the levels of CTL1 and CTL2 expression. **(A)** CTL1 and CTL2 protein levels in Ctrl and M2 cells transfected with CTL1 cDNA, CTL2 cDNA and CTL2 siRNA. **(B)** In M2 cells transfected with CTL1 cDNA, Etn uptake was increased by 25% after 24h and 75% after 48h. In M2 cells transfected with CTL2 cDNA, Etn uptake increased by 25% after 48h. **(C)** Cho uptake at 48h was increased by 50% in Ctrl cells transfected with CTL1 cDNA and 25% in Ctrl cells transfected with CTL2 cDNA. **(D)** Etn uptake was increased by 25% in Ctrl cells transfected with CTL1 cDNA and by 25% in Ctrl cells transfected with CTL2 cDNA. **(E)** [^14^C]-Etn radiolabeling of CDP-Etn pathway and PE synthesis in CTL1 overexpressing M2 cells. **(F)** [^14^C]-Etn 1-3 h pulse-chase analysis of CDP-Etn pathway and PE degradation in CTL1 overexpressing M2 cells. Each bar or point represents the mean ± SD (n = 4), * p < 0.05, ** p < 0.01, *** p < 0.001.

CTL1 cDNA increased Etn transport by 100% and CTL2 cDNA increased Etn transport 25% in M2 cells 48h post transfection (Fig. 4B). In Ctrl cells, CTL1 cDNA increased Cho (Fig. 4C, upper panel) and Etn (Fig. 4D upper panel) transport by 50% and 25% respectively. CTL2 cDNA similarly increased Cho and Etn transport by 25% (Fig. 4C,D-lower panels). Overall, the overexpression data corroborated the kinetic data in Figs 1 and 2 and showed that CTL1 and CTL2 promote transport with similar affinities for Cho and Etn, further showing their functional connection with both arms of the Kennedy pathway.

### Overexpressed CTL1 facilitates CDP-Etn Kennedy pathway in deficient cells

To establish if the overexpression of CTL1 in mutant cells can facilitate CDP-Etn Kennedy pathway, CTL1 cDNA transfected (M2+CTL1) and CTL1 deficient (M2) cells were monitored with [^14^C]Etn radiolabeling in pulse (Fig. 4E) and pulse-chase (Fig. 4F) experiments. As expected, the incorporation of the [^14^C] in Etn, CDP-Etn and PE was significantly increased in CTL1 cDNA transfected relative to untransfected M2 cells. Both types of labelling experiments showed increased P-Etn and CDP-Etn degradation and increased PE synthesis and turnover in M2 transfected cells. Overall, these data demonstrated that the stimulated Etn transport by CTL1 expression increased the flux through the CDP-Etn pathway in CTL1 deficient cells.

### Pharmacological and siRNA inhibition of Etn transport

It is well known that various organic cations can inhibit CTL1 mediated Cho transport (20). We assessed the inhibitory effect of organic cations and CTL2 knockdown on Etn transport in Ctrl and M2 cells (Fig. 5). This helped us understand the magnitude at which CTL1 and CTL2 contributed to Etn transport. Total (CTL1+CTL2) transport (Fig. 5A, Ctrl cells), CTL1 mediated transport (Fig. 5B, Ctrl+CTL2 siRNA cells), CTL2-mediated transport (Fig. 5C, M2 cells) and residual, CTL1 and CTL2 independent transport (Fig. 5D M2 + CTL2 siRNA cells) were quantified. In each case, the inhibitory constants *K*_*i*_ were deduced from semi-log plots (% of remaining Etn transport vs. logM concentration) and compared between different transport conditions.

**Figure 5:**
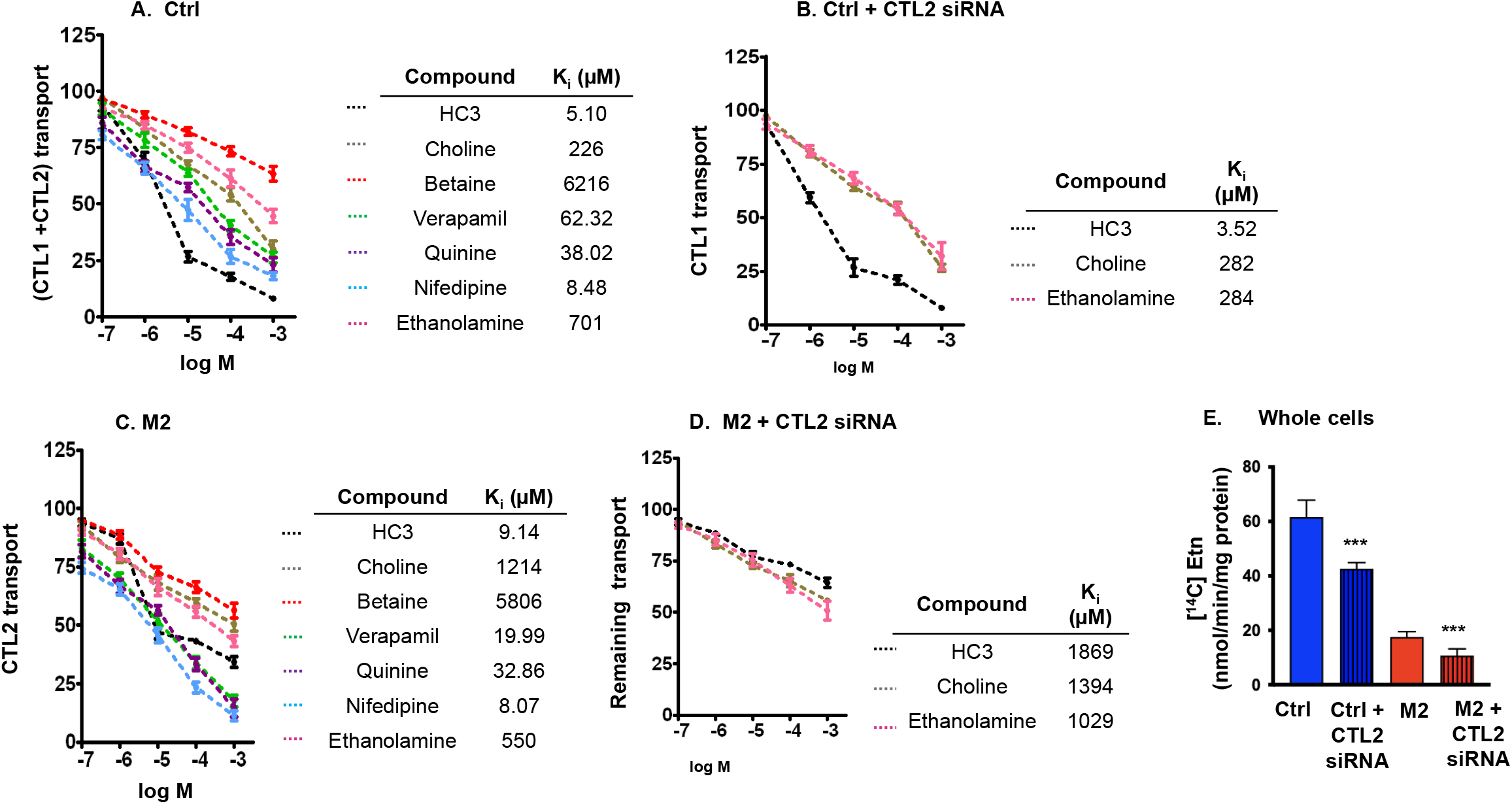
Pharmacological distinction of ethanolamine transport and transporters. **(A)** Semi-log plots of Etn transport inhibition in Ctrl cells and **(B)** Ctrl cells transfected with CTL2 siRNA. The cells were pre-incubated with indicated compounds for 30 minutes and the uptake of 20 μM [^14^C]-Etn was measured for 20 minutes. Semi-log plots of Etn transport inhibition in **(C)** M2 cells and **(D)** M2 cells transfected with CTL2 siRNA. The cells were pre-incubated with test compounds for 30 minutes and the uptake of 20 μM [^14^C]-Etn was measured for 20 minutes. The obtained Ki inhibitory constants for all cell types and treatments are indicated. **(E)** [^14^C]-Etn uptake in the absence of inhibitors in Ctrl and M2 cells with and without siRNA transfection. Each point represents the mean ± SD (n = 4); *** p < 0.001.

As expected, the CTL1 and CTL2 specific inhibitor HC-3 strongly inhibited Etn transport with *K*_*i*_ values of 3.52 μM (CTL1), 9.14 μM (CTL2) and 5.10 μM for the total (CTL1 + CTL2) transport. Nifedipine (a calcium channel blocker), was as potent as HC-3 with *K*_*i*_ = 8.07 - 8.48 μM. Verapamil (a calcium channel blocker) was an intermediate CTL1/2 inhibitor with *K*_*i*_ = 20 - 62 μM. Additionally, quinine (antimalaria drug) with *K*_*i*_ = 33 - 38 μM, was a medium CTL1/2 inhibitor while the Cho oxidation product betaine was a poor inhibitor (*K*_*i*_ = 5806 - 6216 μM) of CTL1 and CTL2 Etn transports (Fig. 5B and C).

The inhibition of CTL1 specific transport (Fig. 5B) with excess choline and Etn was similar, with *K*_*i*_ = 282 - 284 μM. The *K*_*i*_ however differed for CTL2 specific transport (Fig. 5C), with *K*_*i*_ = 1214 μM for Cho and 550 μM for Etn. The *K*_*i*_ value for the total (CTL1 + CTL2) transport (Fig. 5A) was 226 μM for Cho and 701 μM for Etn. Thus, the *K*_*i*_ values indicated that Cho and Etn are transported similarly by the high affinity transporter CTL1. The low affinity transporter CTL2 however had reduced and different affinity for Cho and Etn, with more preference for Etn as its substrate. Residual Etn transport (unrelated to CTL1 and CTL2) was distinguished by CTL2 siRNA treatment of M2 cells (Fig. 5D). This residual transport component has a low affinity for Cho and Etn (*K*_*i*_ = 1029-1394 μM) and three orders of magnitude higher *K*_*i*_ = 1869 μM for HC-3 showing that is not a CTL1/2 related transport. Finally, comparison of all transport velocities (Fig. 5E) showed a general order of contributions, from the high affinity CTL1 (Ctrl + CTL2 siRNA), low affinity CTL2 (M2), and the residual very low affinity (M2+CTL2 siRNA) transports for Etn. CTL1(Ctrl) contributed 80%, CTL2 (M2) 12.5%, and the unrelated residual transport (M2 + CTL2 siRNA) accounted for 7.5% to the total transport. The Ctrl and M2 cells express OCT and OCTN and other unspecific Cho transporters that could be contributing to this residual transport (15).

### CTL1 and CTL2 mediate Etn transport to mitochondria

CTL1 and CTL2 are present in the mitochondria and are involved in mitochondrial Cho transport (4,5). We used COXIV as a marker of mitochondria isolated from Ctrl and M2 cells and CTL2 mRNA expression for the siRNA knockdown of CTL2 transport (Fig. 6A). By comparing the contributions of all mitochondria transport components (Fig. 6B), CTL1+CTL2 (Ctrl) contributed 70%, CTL2 (M2) 20%, and the residual unrelated transport (M2 + CTL2 siRNA) contributed 10% to the total mitochondria Etn transport. We also compared the rates of [^14^C]-Etn transport in the isolated mitochondria and the whole cells and in the presence and absence of the specific inhibitor HC-3. As expected, the CTL1 and CTL2 inhibitor HC-3 blunted Etn uptake in a time dependent manner in the mitochondria (Fig. 6C, E) and the whole cells (Fig. 6 D, F). In the absence of HC-3, the rate of Ctrl mitochondria transport was similar to the whole cell Ctrl transport (0.04 and 0.05 μmol/mg/min, respectively); M2 mitochondrial transport was also similar to the whole cell transport (0.008 and 0.01 μmol/mg/min respectively) (Fig. 6E, F), demonstrating that the same proteins are responsible for the transports in the whole cell and mitochondria. In addition, the M2 mitochondrial (CTL2 only) transport was 5-fold slower that the total (CTL1+CTL2) transport of the Ctrl mitochondria. Taken together, these data established that CTL1 and CTL2 mediate mitochondrial Etn transport with the same kinetic properties as in the whole cells.

**Figure 6:**
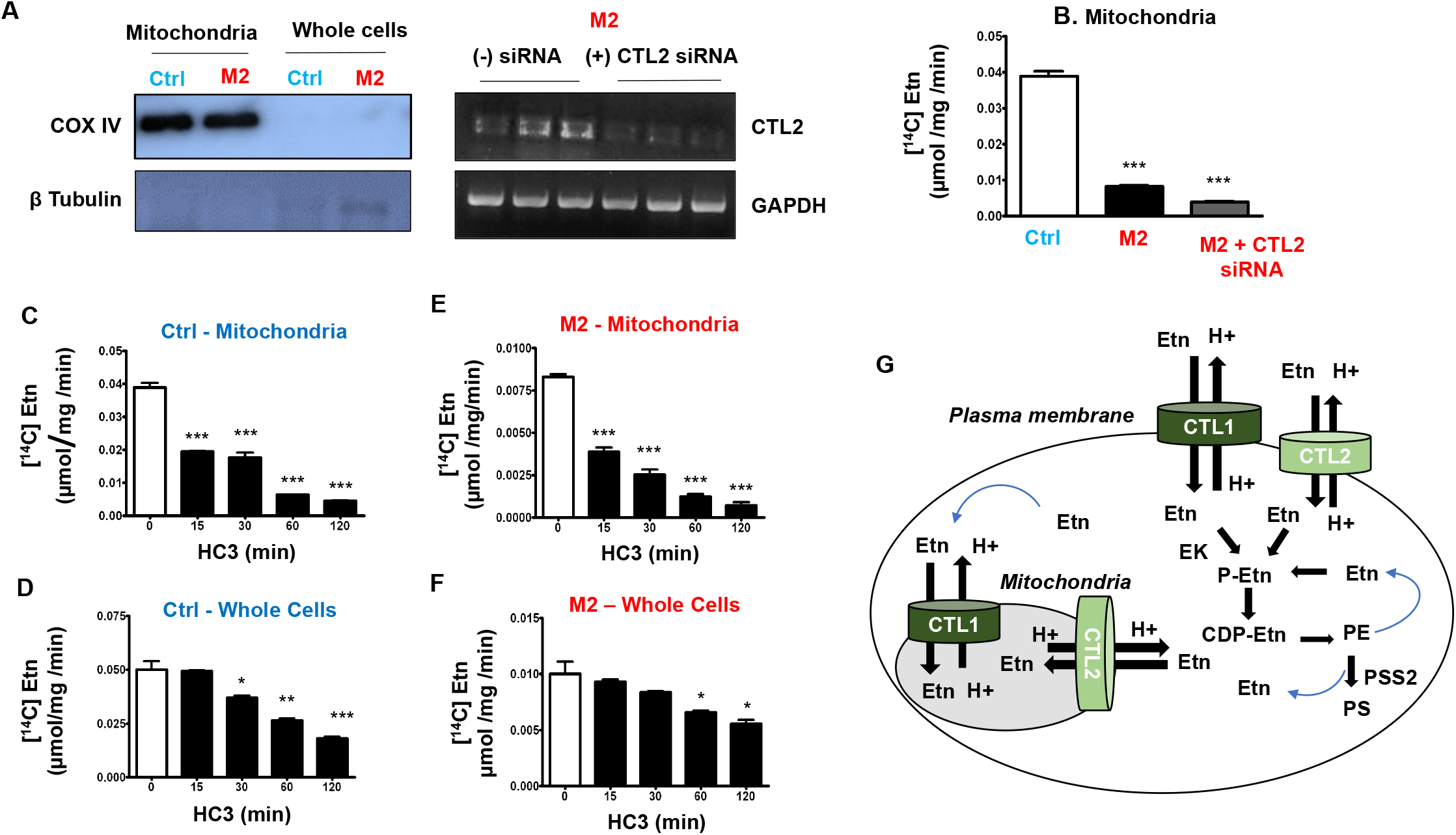
Characteristics of mitochondrial ethanolamine transport. **(A)** Western blots of COXIV mitochondrial marker demonstrating purity of Ctrl and M2 mitochondria; and CTL2 mRNA expression in siRNA-CTL2 treated and untreated M2 cells with complement protein data in Fig. 4A. **(B)** Mitochondrial Etn transport and the effect of CTL2 knockdown in M2 cells; Etn uptake in Ctrl mitochondria was reduced to 25% in M2 (CTL1 deficient) mitochondria and further diminished with siRNA depletion of CTL2. Time course inhibition of Etn uptake in **(C)** Ctrl mitochondria and **(D)** whole cells treated with 20 μM HC-3. Time course of Etn uptake in **(E)** M2 mitochondria and **(F)** M2 cells treated with 20 μM HC-3. Each point or bar represents the mean ± SD (n = 4), * p < 0.05, ** p < 0.01, *** p < 0.001. **(G)** Schematic representation of ethanolamine transport mechanisms at the cell surface and mitochondria. CTL1 and CTL2 are ethanolamine/proton antiporters of high and low affinity, respectively. They are responsible for extracellular uptake and intracellular balance of [Etn]. Extracellular Etn is immediately phosphorylated by ethanolamine kinase (EK) and further consumed by the CDP-Etn pathway to form PE phospholipid. The intracellular Etn is released by PE degradation by lipolysis and PE exchange to PS by PS synthase 2 (PSS2) and it can enter mitochondria by both transporters.

## Discussion

PE and PC are bilayer forming phospholipids involved in fundamental membrane processes, growth, survival and cell signaling (11). PE and PC are similarly synthesized by CDP-Etn and CDP-Cho Kennedy pathway, in which the extracellular substrates Cho and Etn are actively transported into the cell, phosphorylated, and coupled with diacylglycerols (DAG) to form the final phospholipid product. Cho and Etn released from PC and PE also need to be transported in and out of the cytosol and mitochondria or reincorporated into the Kennedy pathway. The plasma membrane CTL1 is firmly assigned to Cho transport for PC synthesis (15), yet the exact function of the mitochondrial CTL1 is still not clear. In the liver and kidney mitochondria, Cho is specifically oxidized to betaine, the major methyl donor in the one-carbon cycle (6). Since broadly expressed, it is proposed that the mitochondrial CTL1 could maintain the intracellular pools of Cho and as ^+^H-antiporter could regulate the electrochemical/proton gradient in the mitochondria (5, 15, 18). CTL2 is only indirectly implicated in Cho transport and until this work the exact CTL2 substrate binding and transport mechanism were not firmly established, in neither the whole cells nor isolated mitochondria.

Based on our extensive work on CTL1 and Cho transport (3-8, 15) and known similarities between Cho and Etn transports in various conditions (12-14) we postulated that CTL1 could be that long-searched for Etn/Cho transporter and the last missing link between CDP-Cho and CDP-Etn pathways for phospholipids synthesis. We conducted an extensive number of kinetic, metabolic, and genetic experiments to solidify this hypothesis. We established that CTL1 mediated a high affinity Etn transport with *K*_*1*_ = 56-67μM and that CTL2 mediated a low Etn affinity transport with *K*_*2*_ = 275-299 μM in primary human fibroblasts and monkey Cos7 cells. Importantly, the CTL1 affinity constant for Etn binding is in the range of physiological Etn concentration in rat and humans (10-75μM) (13), and explains why CTL1 contributed the most (70-80%) of the Etn transport in the whole cells and mitochondria.

We recently described the first human disorder caused by homozygous frame-shift mutations in the CTL1 gene *SLC44A1*: M1= *SLC44A1* ^ΔAsp517^, M2= *SLC44A1* ^ΔSer126^ and M3= *SLC44A1* ^ΔLys90^ (15). After an extensive characterization of transport and metabolism in patient’s fibroblasts it was apparent that diminished Cho transport is the primary cause of this new neurodegenerative disorder with elements of childhood-onset parkinsonism and MPAN (mitochondrial membrane protein-associated neurodegeneration)-like abnormalities. Paradoxically, although Cho transport and CDP-Cho Kennedy pathway were diminished, PC remained preserved in the cerebrospinal fluid and skin fibroblasts of the affected individuals (15). The cell membranes were however drastically remodeled and depleted of PE and PS, apparently as a homeostatic response to preserve PC and prevent Cho deficiency in the affected individuals (15). Since the majority of PE is produced *de novo* by the CDP-Etn pathway, we utilized CTL1 mutant cells to establish if Etn transport and *de novo* PE synthesis were reduced in the affected individuals. We showed that (M2) *SLC44A1* ^ΔSer126^ patient fibroblasts relies on CTL2 for Cho and Etn transport and were devoid of CTL1 mediated Etn transport, which revealed a new biological role for CTL1 and CTL2 that could help in developing new treatments strategy for this devastating disease.

Cho supplementation led the membrane lipids and organelle recovery in CTL1 mutant fibroblast (15) suggesting that transporters other than CTL1 could support the transport when extra substrate became available. We focused on CTL2 as the most plausible alternative candidate for Cho and Etn transport in CTL1 deficient cells (19-22, 25). We separated CTL2 and CTL1 transports using specific antibodies, and depletion and overexpression strategies in CTL1 M2 mutant and Ctrl cells. As a low affinity Etn transporter CTL2 contributed 20-30% to the total transport in the whole cells and isolated mitochondria. CTL1 deficient cells had reduced but not absent CDP-Cho (15) and also had reduced CDP-Etn Kennedy pathway showing that CTL2 is able to channel both substrates for phospholipid synthesis, albeit with a reduced capacity. Apparently, in the individuals with the *SLC44A1* homozygous mutation (15) under normal physiological conditions CTL2 did not compensate the complete loss of CTL1 but as a low affinity (high capacity) transporter it could be beneficial in delivering extra Cho and Etn.

PE plays important structural role in mitochondria (24, 26) and is important for stabilizing the electron transport complexes (27-30). Both CTL1 and CTL2 are prominently present in mitochondria and both transport Cho (3, 19) into mitochondria. When identical proteins are involved, it is expected that transport characteristics are similar even in separate cellular compartments; this was case with Etn transport in the whole cells and mitochondria of Ctrl cells (40-50 nmol/mg/min; CTL1+CTL2) and M2 whole cells and mitochondria (8-10 nmol/mg/min CTL2 only). We clearly demonstrated that CTL1 and CTL2 channel the extracellular Etn into the Kennedy pathway. Why there is a cellular need for active Etn transport and involvement of the low and high affinity transporters in the mitochondria is not clear. Mitochondrial CTL1 and CTL2 could maintain the intracellular pools of Cho and Etn and since they are both proton antiporters, they could be significant regulators of the proton gradient in the mitochondria. Recent studies in CTL2 knockout mice established that that CTL2 mediated mitochondrial Cho transport is critical for ATP and ROS production, platelet activation and thrombosis (31). CTL2 gene *SLC44A2* is well-established the human neutrophil antigen (32), and genetic risk factor for hearing loss, Meniere’s disease and venous thrombosis (33). Neutrophil CTL2 could interact directly with platelets’ integrin α_IIb_β_3_, and induce neutrophil extracellular trap (NETosis) that than promote thrombosis (34). *SLC44A2* knockout mouse is protected against venous thrombosis showing that CTL2 could be an important therapeutic target for the disease (35-37). Our investigation provides insights into the novel function of CTL2/SLC44A2 as an Etn transporter, which will contribute to a better understanding of previous studies and the optimization of prevention and treatment strategies in those various diseases.

Altogether, CTL1 and CTL2 are the physiological Etn transporters in the whole cells and mitochondria, with intrinsic roles in *de novo* PE synthesis by the CDP-Etn Kennedy pathway and intracellular compartmentation of Etn. A schematic representation of Etn transport mechanisms at the cell surface and mitochondria is in Fig 6G. CTL1 and CTL2 are ethanolamine/proton antiporters of high and low affinity, respectively. They are responsible for extracellular uptake and intracellular balance of Etn. Extracellular Etn is immediately phosphorylated by Etn kinase (EK) and further consumed by the CDP-Etn pathway to form the membrane phospholipid PE. The intracellular Etn is released after PE degradation by lipolysis and/or replacement of the Etn headgroup with serine in PE to form PS by PS synthase 2 (PSS2). The metabolically released Etn can be removed, or it can enter mitochondria and/or be phosphorylated and recycled by the CDP-Etn Kennedy pathway.

## Materials and Methods

### Maintenance of cell lines

Control, M1 and M2 primary human skin fibroblasts were maintained in MEM (Fisher Scientific) supplemented with 20% fetal bovine serum and 2% penicillin/streptomycin. Cells were kept in a humidified atmosphere at 37°C and 5% CO_2_. MCF-7 human breast cancer cells, MCF-10 human mammary epithelial cells and COS-7 fibroblast-like monkey cells were maintained in DMEM (Fisher Scientific) supplemented with 10% fetal bovine serum and 2% penicillin/streptomycin.

### Transport Studies

According to our previously standardized protocols (3) cells were incubated with 0.2 μCi [^14^C]-Etn or [^3^H]-Cho with the compound of interest for 20 minutes and room temperature. To stop the transport, cells were washed in ice cold KRH buffer containing 500 μM ‘cold’ Cho or Etn. Cells were then lysed in 500 μl ice cold lysis buffer (10 mM Tris-HCl, 1 mM EDTA and 10 mM NaF) and the radiolabeled Cho and Etn were analyzed by LSC. Kinetic constants for Etn transport were determined as before for Cho (38). Cells were incubated (20 min) with increasing concentrations of unlabeled Etn (0-1000 μM) before being treated with 0.2 μCi [^14^C]-Etn for 20 minutes. The saturation curves of [^14^C]-Etn transport velocity (V) vs. Etn concentration (S) were produced in different cell types and the transport affinity constants (K1 and K2) derived from linearized Eadie-Hofstee plots using GraphPad Prism software (GraphPad, Inc.). To study the effect of pH on Etn uptake, the cells were treated with KRH buffers of varying pH (pH 5.5 - 8.5). Buffers were prepared by mixing 10 mM MES (pH 5.5) and 10 mM bicine (pH 8.5). To assess the effects of [Na^+^] on Etn uptake, the cells were subjected to either standard KRH buffer or KRH buffer with LiCl instead of NaCl. For transport inhibition, 8.0 x 10^4^ cells/well were treated with various compounds for 24h. The concentration curves for a specific compound (HC3, Eth, Cho, betaine, verapamil, quinine, nifedipine) were produced for each cell type and Ki derived from the semi log plots of transport Inhibition (%) vs log [drug concentration] were compared. In studies with CTL1 and CTL2 antibodies 8.0 x 10^4^ cells/well were seeded in 6-well plates and grown for 24h. After 24h of growth, various amounts of antibodies were added to cells and incubated for 24h and Cho and Etn uptake were then conducted.

### Radiolabeling of the CDP-Etn Kennedy pathway and phospholipids

To analyze the CDP-Etn Kennedy pathways cells were radiolabeled with 0.2 μCi [^14^C]-Etn (ARC St. Louis, MO) for 1-3h (pulse and pulse-chase). Water-soluble pathway intermediates (Etn, P-Etn and CDP-Etn) and PE were extracted and separated with the method of Bligh and Dyer (39). For pulse experiments, cells were incubated for 1, 2 and 3h with 0.2 μCi [^14^C]-Etn, and for pulse-chase experiments, cells were incubated for 1h with 0.2 μCi [^14^C]-Etn, washed with PBS and then chased for 1-3h with an excess of unlabeled Etn. At each time point, radiolabeled compounds were extracted with the method of Bligh and Byer, separated by TLC using appropriate standards and radioactivity (dpm) determined by liquid scintillation counting (40). Phospholipid (PC, PS and PE) and neutral lipid (DAG and TAG) pools were determined by 24h steady-state (equilibrium) radiolabeling with 3H-glycerol as previously described (40).

### Pcyt2 Activity Assay

The assay was conducted as described (40). In brief, cells were cultured for 24h and 50 μg protein was dissolved in Pcyt2 reaction mixture. The mixture was treated for 15 minutes with 0.2 μCi [^14^C]-PEtn and the reaction was terminated by boiling for 2 minutes. The radiolabeled Pcyt2 reaction product [^14^C]-CDP-Etn was isolated by TLC and Pcyt2 activity was expressed as nmol/min/mg protein.

### RNA extraction and RT-PCR

Total RNA was isolated with TRIzol (Invitrogen, Life Technologies Incorporated, Burlington, ON, Canada). DNase I was used to eliminate genomic DNA and cDNA was synthesized from 2 μg RNA with RNA SuperScript III Reverse Transcriptase (Invitrogen, Life Technologies Incorporated). Expression of CTL1, CTL2, CK, EK, Pcyt1, Pcyt2, PSD, PSS1 and PSS2 was determined by PCR using the primers and conditions as before (15). Reactions were standardized by amplifying glyceraldehyde 3-phosphate dehydrogenase (GAPDH) and relative band intensity was quantified using ImageJ software (NIH, Bethesda, MD, USA).

### Expression of rat CTL1-Myc cDNA, murine CTL2 cDNA and CTL2 siRNA

Confluent cells were transfected with 5 μg of pCMV3-ORF-C-Myc-SLC44A1 cDNA (Sino Biologics, # RG80408-CM), pCMV-SPORT6-SLC44A2 cDNA (Genomics Online, # ABIN3822596) or empty vector (pcDNA4His-Max B, # V86420) using Lipofectamine 2000 (Invitrogen). Transfections with 30 nM SLC44A2 siRNA (Santa Cruz Biotechnology, # sc-62163) was with siPORT lipid transfection agent.

### Immunoblotting

Cells were washed 3x with PBS and subjected to a lysis buffer (25 mM Tris, 15% glycerol, 1% Triton X-100, 8 mM MgCl_2_, 1 mM DTT, protease inhibitor cocktail and phosphatase inhibitor cocktail) at 4°C for 30 minutes. Protein concentration was determined with the bicinchoninic (BCA) assay (Pierce, Rockford, IL, USA). The LV58 (N-terminus) and ENS-627 (C-terminus) antibodies (both 1:500 in 5% skim milk in TBS-T) detect the 72 kDa CTL1 protein under native (non-denaturing) conditions. Samples were mixed with loading buffer (62 mM Tris-HCl, 0.01% bromophenol blue and 10% glycerol) and separated by PAGE at 120 V for 1.5h. CTL1, CTL2 (Abnova; 1:200 in 5% skim milk in TBS-T) and β-tubulin (Cell signaling; 1:1000 in 5% skim milk in TBS-T) were resolved on an 8% native gel and proteins were transferred onto PVDF membranes (Pall Canada, Mississauga, ON, Canada) by a semi-dry transfer system and stained with Ponceau S. Membranes were blocked in 5% skim milk in Tris Buffered Saline-Tween 20 (TBS-T) solution and then incubated with primary antibodies (1:500 in 5% skim milk in TBS-T) overnight at 4°C. Membranes were washed with TBS-T and then incubated with an anti-rabbit horseradish peroxidase conjugated secondary antibody (New England Biolabs, 1:10,000 in 5% skim milk in TBS-T) for 2h. Membranes were washed in TBS-T and proteins were visualized using a chemiluminescent substrate (Sigma Aldrich, Oakville, ON, Canada).

### Mitochondrial Isolation

Mitochondria were isolated as initially described (18). In brief, cells were incubated for 20 minutes in ice-cold RSB swelling buffer and homogenized; 19 ml MS buffer was added, and the cell homogenate was centrifuged at 2500 rpm for 5 minutes. This step was repeated twice, and the final supernatant was centrifuged at 12,500 rpm. The resulting pellet was resuspended in MS buffer. The mitochondria purity was determined using COXIV mitochondria marked and β-tubulin as a whole cell control.

### Statistical Analysis

All measurements are expressed as means from quadruplets ± SEM. Statistical analysis was performed using GraphPad Prism software (GraphPad, Inc.). Data were subjected to students T-test. Differences were considered statistically significant at *p < 0.05.

## Data Availability

All study data are included in the article

## Author contributions

A. T. and M.B. designed research; A.T. J. I., and S.G. performed research; A.T and M.B. analyzed data and wrote the paper.

The authors declare no competing interests.

## Acknowledgments

We thank Christina Fagerberg (Odense University, Denmark) and Felix Distelmaier (Heinrich-Heine University, Dusseldorf, Germany) for CTL1 mutant fibroblasts. This work was supported by the University of Guelph Scholarship (to A.T. and S.G.), the Canadian Institutes of Health Research Grant # CIHR-450137 and the National Sciences and Engineering Research Council of Canada Discovery Grant # NSERC-400482 (to M.B.)

